# Rapid tRNA Isolation and Chemiluminescent Northern Blot Detection of tRNA and tRNA-Derived Fragments

**DOI:** 10.1101/2025.10.14.681074

**Authors:** Pavlina Gregorova, Minna-Maria K. Heinonen, Milla M. Laarne, L. Peter Sarin

## Abstract

Transfer RNA (tRNA), its post-transcriptional modifications, and tRNA-derived fragments (tRFs) play essential roles in cellular processes and gene regulation. Here, we present a fast and efficient tRNA isolation using silica spin columns. To analyze the isolated tRNA and detect tRFs, we describe a sensitive and cost-effective non-radioactive Northern blotting technique. Additionally, this blotting method is compatible with chemical affinity modifiers, such as [p-(N-acrylamino)-phenyl]mercuric chloride (APM) or 3-(acrylamido)phenylboronic acid (APB) enabling the detection of chemical modifications in specific tRNA isoacceptors.

**Graphical abstract:** 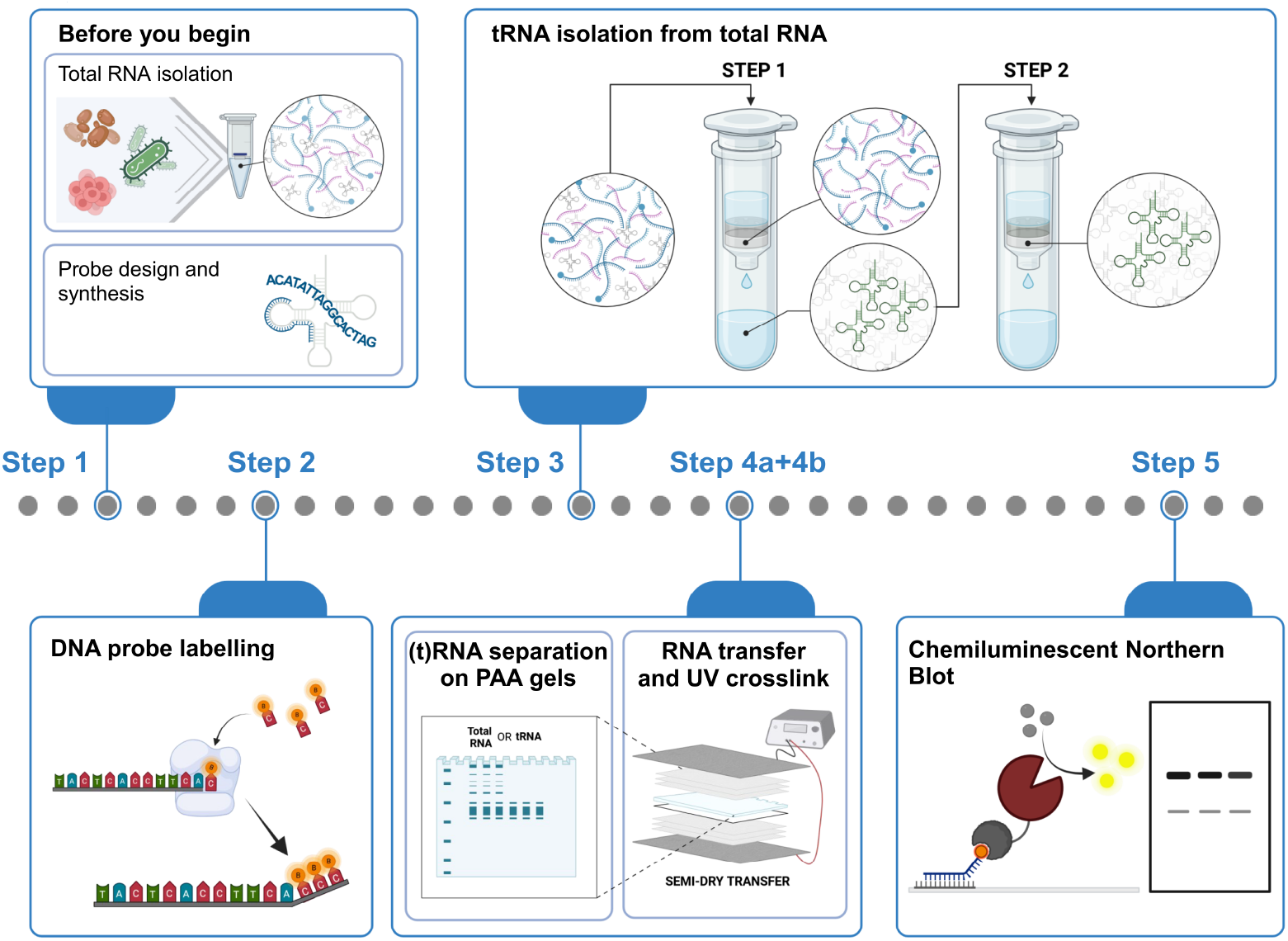

**Highlights:**

- Rapid (<30 min) tRNA isolation from total RNA using silica containing spin columns
- Procedure for DNA probes biotinylation via terminal deoxynucleotidyl transferase
- Non-radioactive chemiluminescent Northern blotting

## Before you begin

The tRNA isolation method presented here requires total RNA of good quality and purity. For total RNA extraction, various extraction techniques can be employed depending on the organism of interest. Ensure that the initial total RNA does not contain contaminants, such as lipids. Although the tRNA isolation method is broadly applicable to total RNA extracted from any organism, this protocol demonstrates tRNA isolation from bacteria and yeast. The isolated tRNA is suitable for various downstream applications, including sequencing and mass spectrometry analyses.

The design of probes is critical for achieving specific and accurate Northern blot detection. Enzymatic probe labeling facilitates the testing of multiple probe designs without the need for costly commercial labeling. When designing probes for tRNA detection, it is advisable to consult modification data from resources such as MODOMICS^1^ to account for possible tRNA modifications that may interfere with probe hybridization. Once the newly designed probes are validated, commercial biotin-labeled versions can be obtained for long-term use. In this protocol, three validated probes are presented: two targeting tRNA-Leu-UAG and tRNA-His-GUG from the gram-negative bacterium *Shewanella glacialimarina* TZS-4T^2^, and one targeting tRNA-Glu-UUC from *Saccharomyces cerevisiae* BY4741.

## Innovation

Existing tRNA purification protocols are predominantly based on anion-exchange chromatography, which utilizes selective tRNA binding to quaternary amino groups in the presence of sodium ions^2-4^. Even though such techniques yield tRNA of sufficient purity, they are laborious and require extensive handling times, in addition to which the purified tRNA often elutes into multiple fractions, necessitating additional precipitation steps^3^. Furthermore, these approaches may also use FPLC or HPLC equipment, which further limits their applicability. In contrast, this protocol describes an optimized silica spin column tRNA isolation method, which enables simple and fast isolation while minimizing undesired rRNA contamination. The resulting tRNA is ready to use for downstream analyses, such as northern blotting or RNA mass spectrometry.

While significant advancements have been made in chemiluminescent blotting methods^5-7^, the existing methods typically use commercial probes with a single biotin label. In this protocol, the detection sensitivity is substantially enhanced by incorporating multiple biotin-carrying nucleotides using a terminal deoxynucleotidyl transferase (TdT), resulting in improved detection of targets with low abundance, such as single tRNA isoacceptors and tRNA-derived fragments.

### Probe design and synthesis

**Timing: 1h-5 days (depending on DNA synthesis service supplier)**

1. Design the DNA oligonucleotide probe for detection of desired tRNA.
  a. For a successful design, carefully considered key factors, such as probe sequence length, GC content, and sequence specificity. The probes used with the hybridization buffer defined in this protocol should have the following properties:
    i. Melting temperature: 65 to 85 °C
    ii. GC content: 45-80%
    iii. Length: 28 to 35 nt ***Note***: Calculate Tm of each probe using IDT’s OligoAnalyzer tool (https://www.idtdna.com/calc/analyzer) using following parameters: 500 mM Na^+^, 0.0001 μM oligo conc., 0 mM Mg^2+^, 0 mM dNTPs conc.
2. Synthesize the DNA probes using your DNA synthesis service of choice
  a. Desalting provides sufficient probe purity for labeling.
3. Upon arrival, dissolve DNA probes in RNase-free water to 100 μM concentration. Store at −20°C.

### Total RNA isolation

**Timing: 1-2 days (depending on organism)**

4. Grow cells in an appropriate medium and collect as recommended.
5. Isolate total RNA using a method suitable for the organism of interest.
  a. For gram negative bacteria, such as *Shewanella glacialimarina* TZS-4T, use TRIzol (or other equivalent reagent) followed by acidic phenol re-extraction as described in ref. ^2^
  b. For yeast, use acidic phenol isolation method as described in ref. ^4^ ***Note***: Yeast total RNA isolated by acidic phenol method described in ref.^4^ is enriched for tRNA and will result in higher yields after tRNA purification. ***Note***: Total RNA can be also isolated by commercial column-based kits. However, make sure that the kit isolates RNAs longer than 20 nt. ***CRITICAL***: When isolating total RNA from tissue, ensure that total RNA samples do not contain lipids, as their presence will interfere with RNA binding to the silica column^8^ during tRNA isolation, resulting in a decreased tRNA yield.
6. Dissolve total RNA in RNase-free water and determine the concentration by measuring absorbance at 260 nm or by fluorescent methods (e.g. Qubit). Store at −80 °C.

## Key resources table

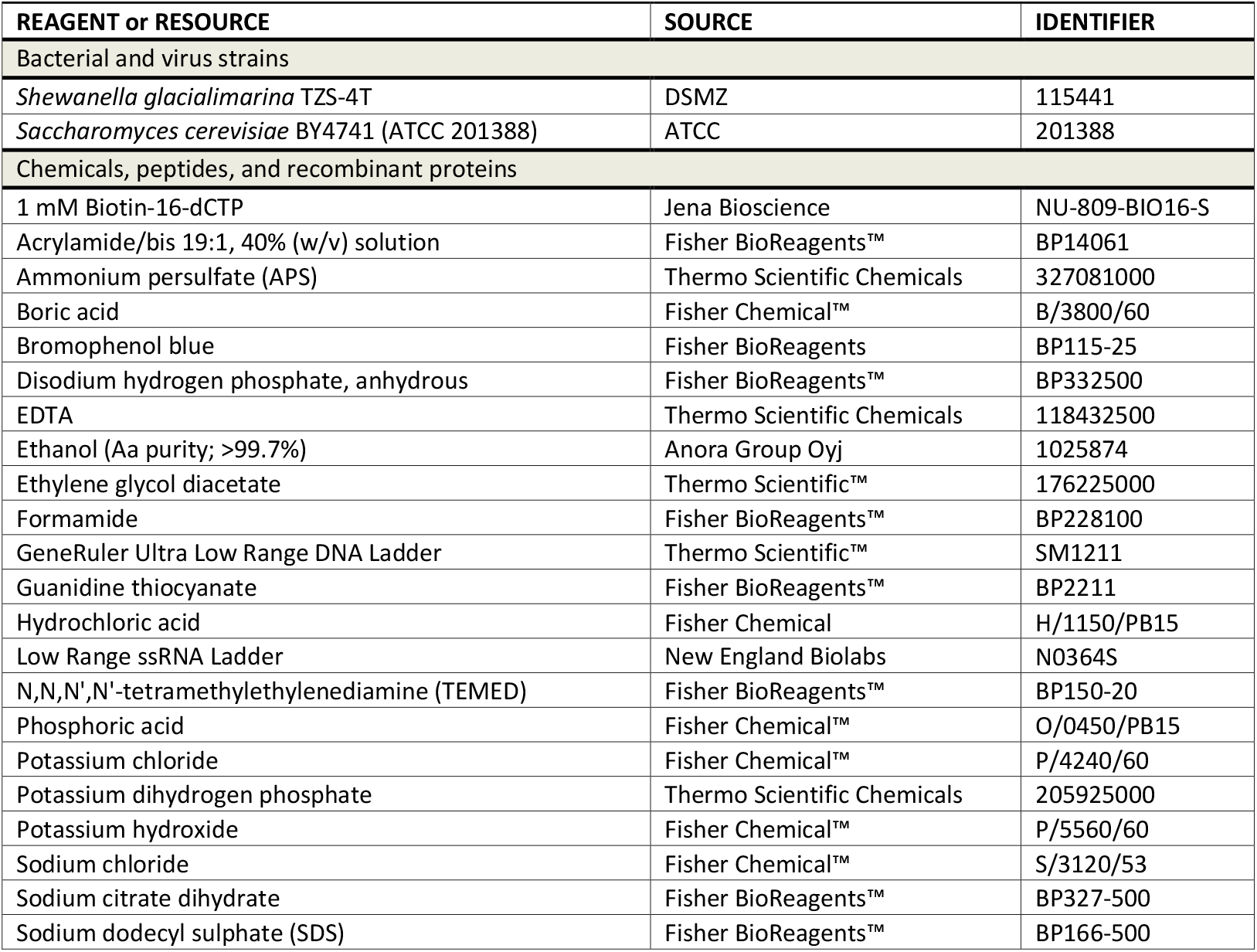
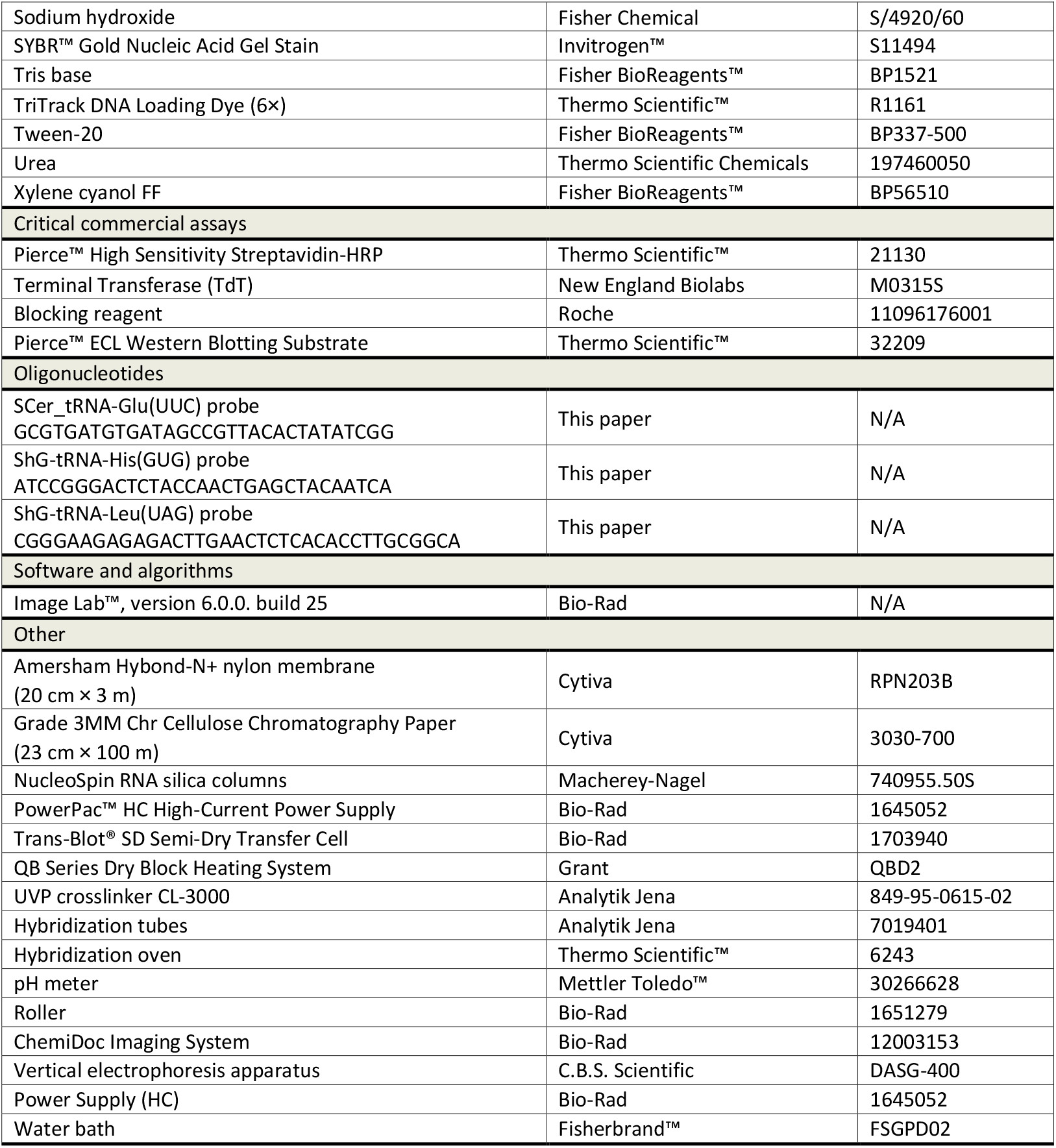

## Materials and equipment setup

**Long RNA binding buffer (LBB)**

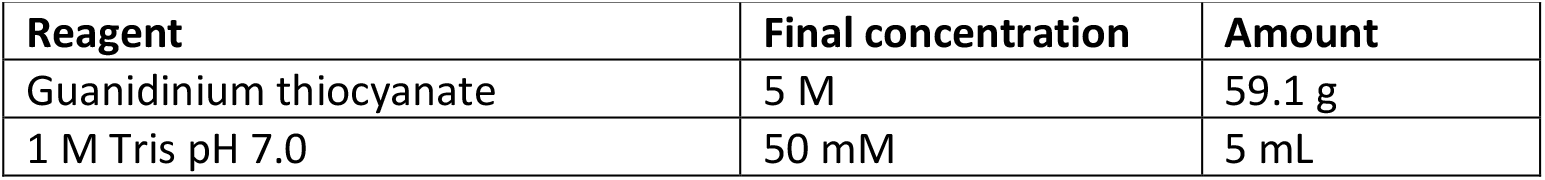
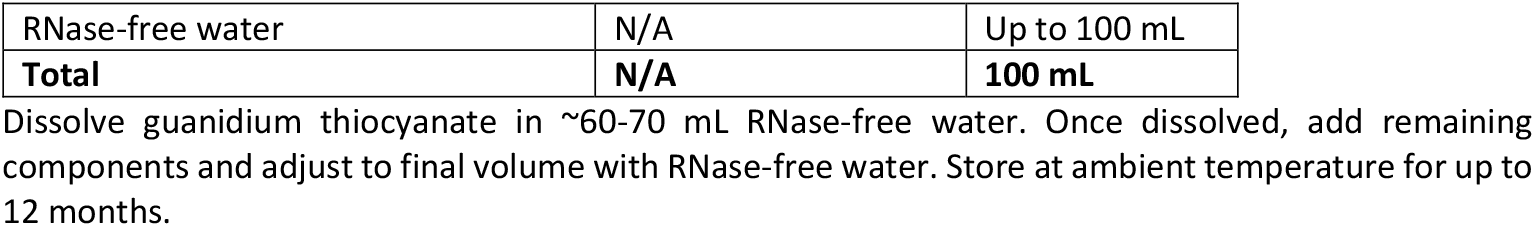

Materials and equipment setup Long RNA binding buffer (LBB)
| Reagent | Final concentration | Amount |
| --- | --- | --- |
| Guanidinium thiocyanate | 5 M | 59.1 g |
| 1 M Tris pH 7.0 | 50 mM | 5 mL |
| RNase-free water | N/A | Up to 100 mL |
| **Total** | **N/A** | **100 mL** |

Dissolve guanidium thiocyanate in ~60-70 mL RNase-free water. Once dissolved, add remaining components and adjust to final volume with RNase-free water. Store at ambient temperature for up to 12 months.

***CRITICAL***: Do not use if the buffer precipitated during storage.

**Short RNA binding buffer (SBB)**

| Reagent | Final concentration | Amount |
| --- | --- | --- |
| Ethylene glycol diacetate | 80% | 800 µL |
| LBB<br>(5 M Guanidinium thiocyanate,<br>50 mM Tris pH 7.0) | 1 M | 200 µL |
| **Total** | **N/A** | **1 mL** |

Prepare freshly before use.

***Note***: The SBB buffer is conveniently prepared by mixing LBB with EGTA in 1:4 ratio (e.g. 200 μL LBB and 800 μL of EGTA). Isolation of one sample requires ∼700 μL.

**Chaotropic wash buffer (CWB)**

| Reagent | Final concentration | Amount |
| --- | --- | --- |
| 5M Guanidine thiocyanate | 1.6 M | 32 mL |
| Ethanol (pure) | 66% | 66 mL |
| 1M Tris, pH 7.0 | 20 mM | 2 mL |
| **Total** | **N/A** | **100 mL** |

Mix all components together and store at ambient temperature for up to 12 months.

**Ethanol wash buffer (EWB)**

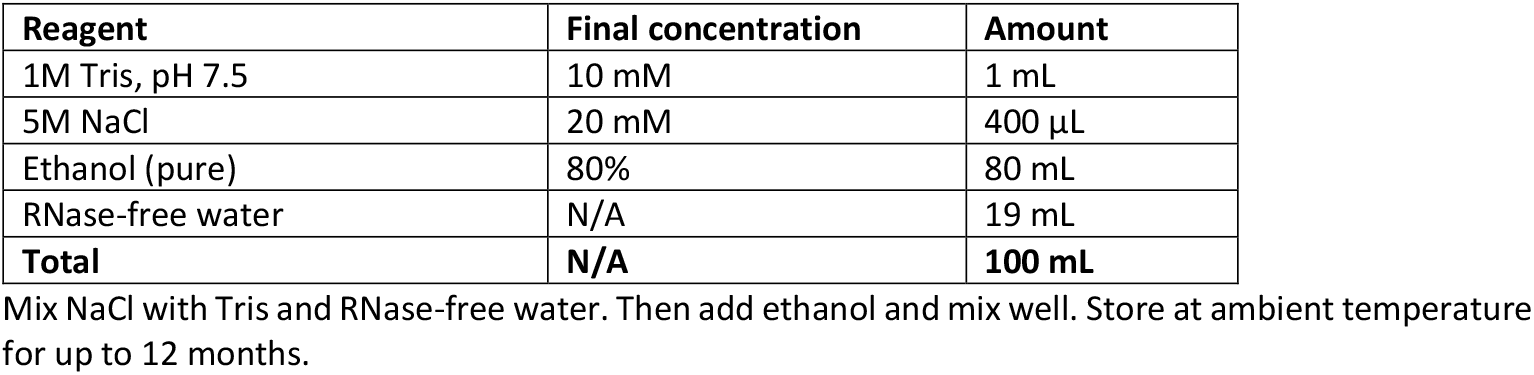

| Reagent | Final concentration | Amount |
| --- | --- | --- |
| 1M Tris, pH 7.5 | 10 mM | 1 mL |
| 5M NaCl | 20 mM | 400 µL |
| Ethanol (pure) | 80% | 80 mL |
| RNase-free water | N/A | 19 mL |
| **Total** | **N/A** | **100 mL** |

Mix NaCl with Tris and RNase-free water. Then add ethanol and mix well. Store at ambient temperature for up to 12 months.

***Note***: Adding concentrated NaCl directly to ethanol will lead to NaCl precipitation. The precipitate dissolves quickly after addition of water. In case of precipitation, mix solution well until precipitate disappears. This does not impact the functionality of the solution.

**10× PBS**

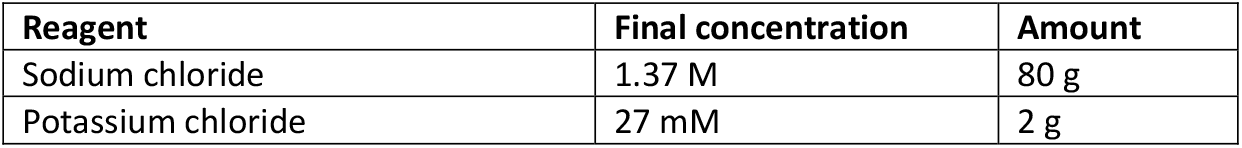
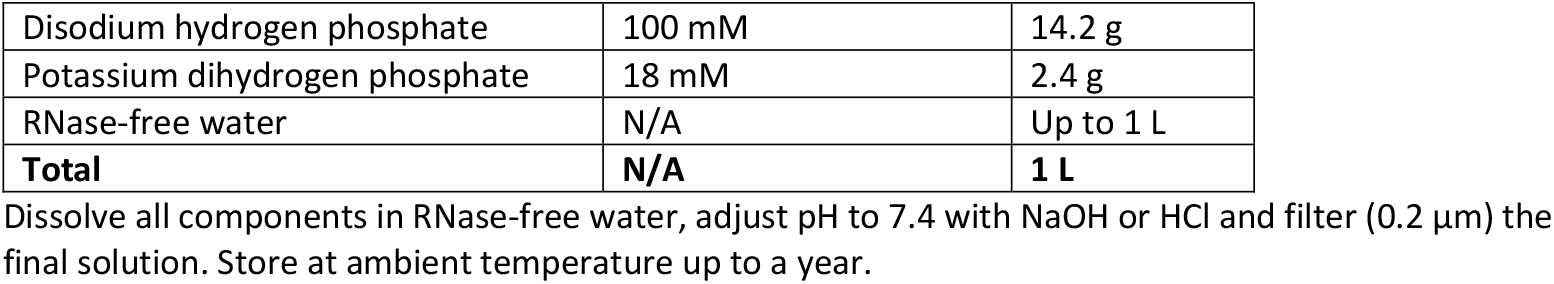

**20× SSC**

| Reagent | Final concentration | Amount |
| --- | --- | --- |
| Sodium chloride | 3 M | 175.3 g |
| Trisodium citrate | 0.3 M | 88.3 g |
| RNase-free water | N/A | Up to 1 L |
| **Total** | **N/A** | **1 L** |

**5× TBE**

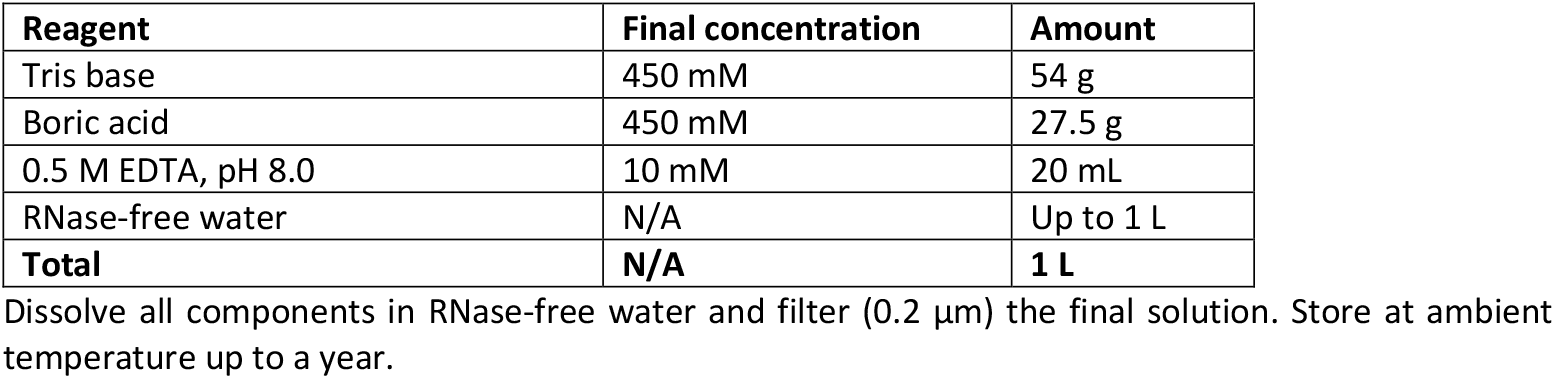

**2× RNA loading dye**

| Reagent | Final concentration | Amount |
| --- | --- | --- |
| Formamide | 90% (v/v) | 9 mL |
| 5× TBE | 0.5× | 1 mL |
| 20% SDS | 0.05% | 25 µL |
| Bromophenol blue | 0.025% | 2.5 mg |
| Xylene cyanol FF | 0.01% | 1 mg |
| **Total** | **N/A** | **10 mL** |

Store at ambient temperature for up to three months or at -20 °C for up to two years.

**Native PAA gel, 12%**

| Reagent | Final concentration | Amount |
| --- | --- | --- |
| 5× TBE | 1× | 4 mL |
| Acrylamide/bis 19:1, 40% (w/v) solution | 12% | 6 mL |
| 10% APS | 0.1% | 200 µL |
| TEMED | 0.1% | 20 µL |
| RNase-free water | N/A | 10 mL |
| **Total** | **N/A** | **20 mL** |

Prepare freshly before use.

**Denaturing PAA Gel Buffer A**

| Reagent | Final concentration | Amount |
| --- | --- | --- |
| 5× TBE | 0.5× | 20 mL |
| Urea | 7M | 84.08 g |
| RNase-free water | N/A | Up to 200 mL |
| **Total** | **N/A** | **200 mL** |

**Denaturing PAA Gel Buffer B**

| Reagent | Final concentration | Amount |
| --- | --- | --- |
| 5× TBE | 0.5× | 20 mL |
| Acrylamide/bis 19:1, 40% (w/v) solution | 20% | 100 mL |
| Urea | 7M | 84.08 g |
| RNase-free water | N/A | Up to 200 mL |
| **Total** | **N/A** | **200 mL** |

**Denaturing PAA gel, 10%**

| Reagent | Final concentration | Amount |
| --- | --- | --- |
| Buffer A | N/A | 10 mL |
| Buffer B | 10% (acrylamide) | 10 mL |
| TEMED | 0.1% | 20 µL |
| 10% APS | 0.1% | 200 µL |
| **Total** | **N/A** | **20 mL** |

Prepare freshly, just before use.

**0.5M phosphate buffer, pH 7.2**

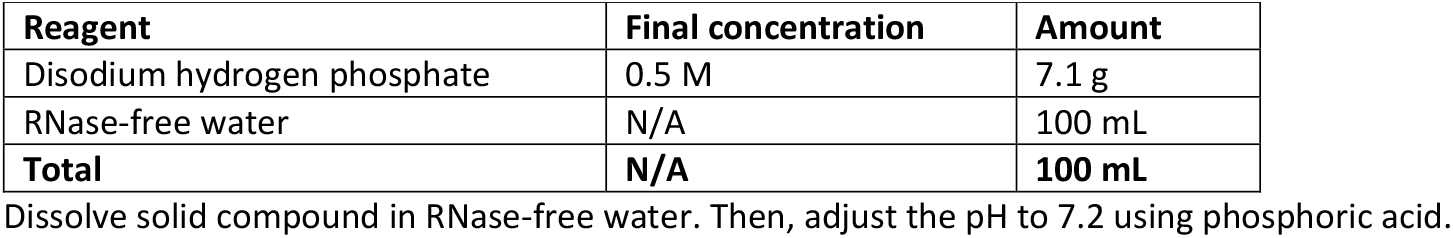

| Reagent | Final concentration | Amount |
| --- | --- | --- |
| Disodium hydrogen phosphate | 0.5 M | 7.1 g |
| RNase-free water | N/A | 100 mL |
| **Total** | **N/A** | **100 mL** |

Dissolve solid compound in RNase-free water. Then, adjust the pH to 7.2 using phosphoric acid.

***CRITICAL:*** pH adjustment takes only a few mL of 50% phosphoric acid. Do not use concentrated acid! Filter (0.2 μm) and store at RT.

**5% Roche blocking reagent (RBR) stock**

| Reagent | Final concentration | Amount |
| --- | --- | --- |
| Roche blocking reagent | 5% | 5 g |
| 0.05 M phosphate buffer, pH 7.2 | 0.05 M | 100 mL |
| **Total** | **N/A** | **100 mL** |

Dissolve 5 g in ~80 mL of 0.05 M phosphate buffer, pH 7.2 by heating it in a microwave oven. The 0.05 M phosphate buffer is prepared by diluting 0.5 M phosphate buffer, pH 7.2 with RNase-free water. Make sure the reagent is properly dissolved. Cool down, adjust to 100 mL and aliquot. Store at -20 °C up to a year.

***CRITICAL:*** Do not boil the solution as it will lead to irreversible precipitation of the blocking reagent.

**Pre-/hybridization buffer**

| Reagent | Final concentration | Amount |
| --- | --- | --- |
| 0.5 M phosphate buffer, pH 7.2 | 0.25 M | 50 mL |
| 0.5 M EDTA | 1 mM | 0.2 mL |
| 20% SDS | 10% | 50 mL |
| Roche blocking reagent | 0.5% | 0.5 g |
| **Total** | **N/A** | **100 mL** |

Dissolve RBR in 0.5M phosphate buffer by heating it in a microwave oven. When all BBR is properly dissolved let the bottle cool down (~ambient to 45 °C). Then, add rest of the components and mix well. Use directly or let cool down closer ambient temperature before storing at 4 °C.

***Note***: Hybridization buffer will heavily precipitate at 4 °C. Prior to use, warm up the buffer at 65 °C in a water bath to dissolve it completely. This step can take up to 1h. The hybridization buffer can be heated and cooled multiple times.

**Membrane wash buffer**

| Reagent | Final concentration | Amount |
| --- | --- | --- |
| 20× SSC | 2× | 100 mL |
| 20% SDS | 0.1% | 5 mL |
| RNase-free water | N/A | Up to 1 L |
| **Total** | **N/A** | **1 L** |

**Blocking buffer**

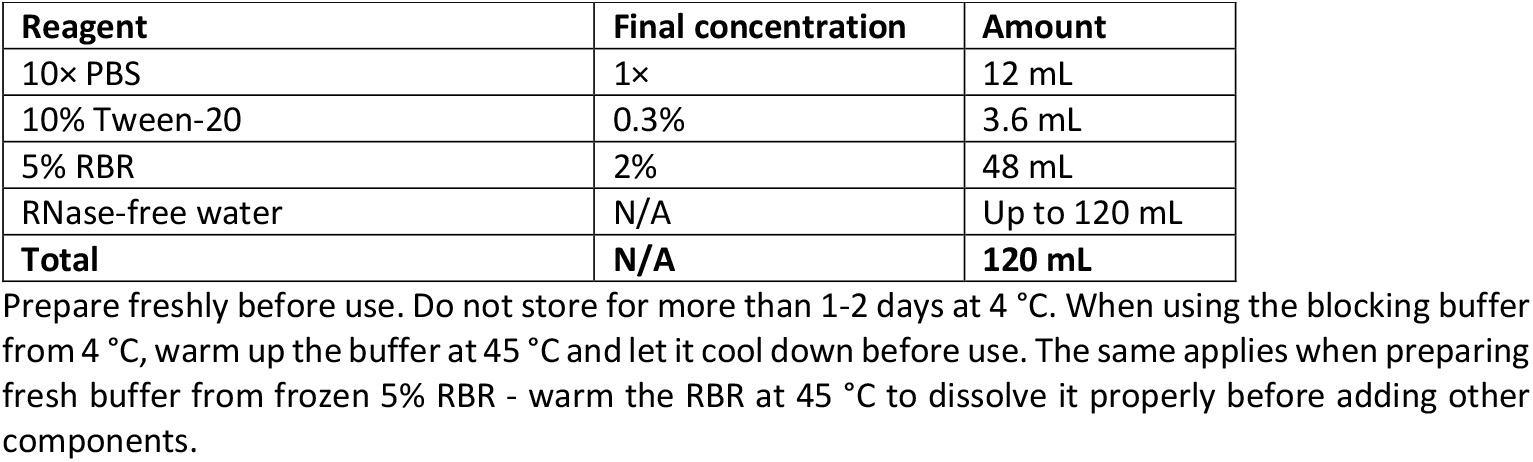

***CRITICAL***: Do not use blocking buffer if it contains any precipitate. This will result in inefficient blocking and higher background!

**PBS wash buffer**

| Reagent | Final concentration | Amount |
| --- | --- | --- |
| 10× PBS | 1× | 100 mL |
| 10% Tween-20 | 0.3% | 30 mL |
| RNase-free water | N/A | Up to 1 L |
| **Total** | **N/A** | **1 L** |

**Detection buffer**

| Reagent | Final concentration | Amount |
| --- | --- | --- |
| Potassium dihydrogen phosphate | 0.05 M | 6.8 g |
| RNase-free water | N/A | Up to 1 L |
| **Total** | **N/A** | **1 L** |

***CRITICAL*:** To minimize RNase contamination, adhere to standard RNA handling procedures throughout the protocol. Use RNase-free plasticware and glassware (baked at 180 – 220 °C for several hours).

***CRITICAL*:** Acrylamide/bis solutions, guanidine thiocyanate, TRIzol, formamide and SDS are harmful. Follow appropriate protective measures and institutional guidelines for handling and disposal.

***Alternatives***: Gel electrophoresis system can be substituted by any other vertical electrophoresis system (e.g. Mini-PROTEAN Tetra Cell, Bio-Rad) which uses smaller (mini) gels. This enables the use of commercial pre-cast gels, such as Criterion TBE-Urea Precast Gels, Bio-Rad.

***Alternatives***: DNA probes can be ordered either 3’ or 5’ pre-biotinylated.

***Alternatives***: Commercial denaturing 2× RNA loading dyes can be used (e.g. RNA Gel Loading Dye (2×), Thermo Scientific, R0641).

***Alternatives***: More sensitive chemiluminescent substrate can be used (e.g. SuperSignal West Pico PLUS, Thermo Scientific, 34580) to enhance the signal detection.

## Step-by-step method details

### tRNA isolation from total RNA

**Timing: 20-60 min (depending on number of samples)**

This section describes the two-step tRNA isolation method, whereby RNA molecules shorter than 120 nt are selectively recovered from total RNA (Figure 1A). Optionally, this isolation process may also be applied as a one-step method to obtain RNA molecules longer than 120 nt.

**Figure 1:**
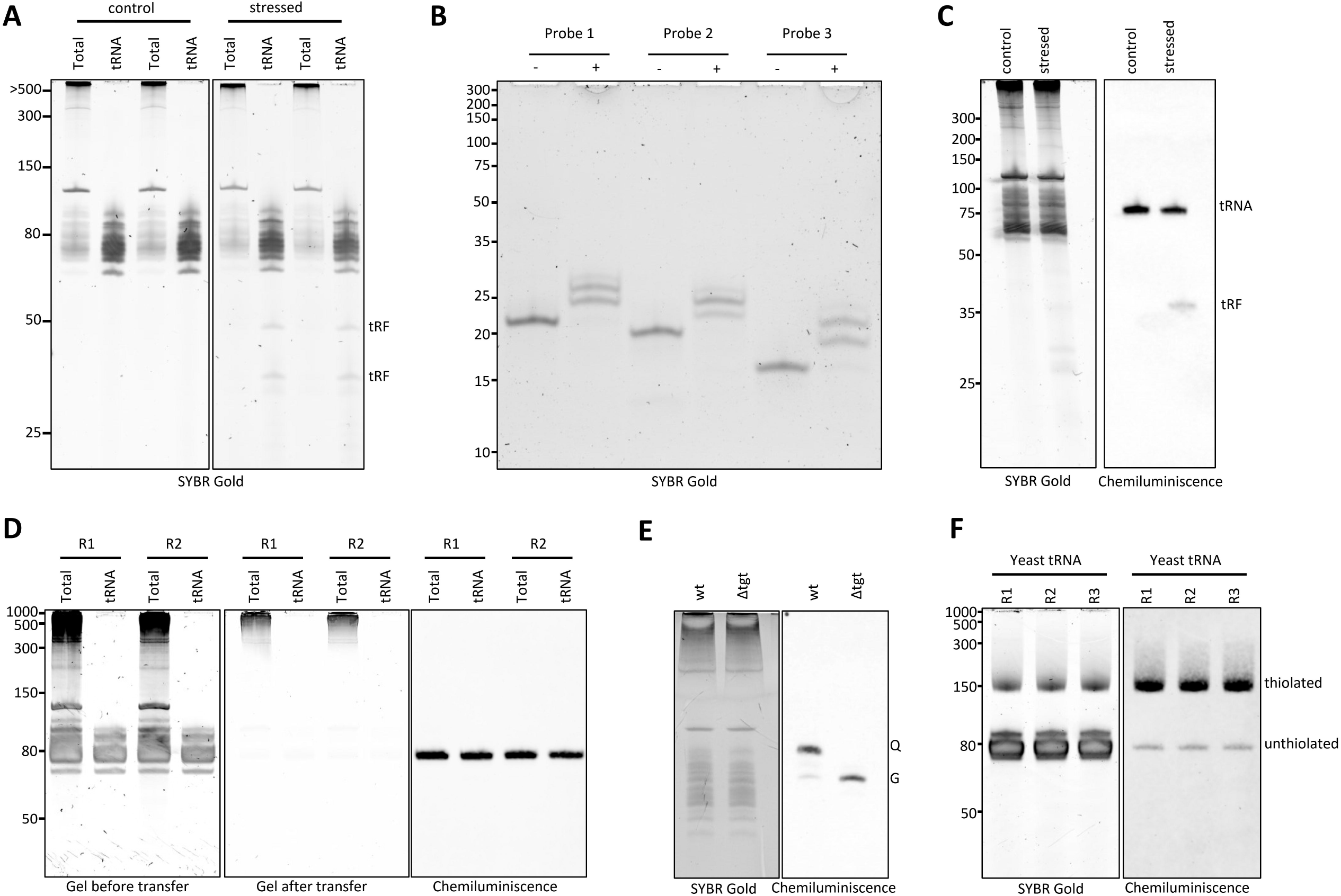
tRNA isolation, quality control steps and expected outcomes of the Northern blot protocol. **A)** Total RNA and tRNA (100 ng for each lane) analyzed on 10% urea-PAA gel to verify purity and quality of isolated tRNA. Note the enrichment of tRNA fragments (tRFs) in the tRNA fractions compared to total RNA. Numbers on the left correspond to the Low Range ssRNA Ladder. **B)** Native 12% PAA gel analysis of 0.4 pmol of unlabeled (-) and labeled (+) probes confirming successful biotinylation. Ladder - Ultra-low range DNA ladder. **C)** Analysis of bacterial total RNA (2 μg) by gel electrophoresis (left) and detection of tRNA-Pro and its stress-induced fragments (tRF; right) using long (14x 10 cm) 12% urea-PAA gels. Ladder - Low Range ssRNA Ladder. **D)** Typical Northern blot detection of bacterial tRNA-His. Total RNA (2 μg) and tRNA (200 ng) are separated on 10% urea-PAA mini (8.6 × 6.7 cm) gels (left), followed by verification of transfer (middle) and finally, tRNA-His is detected by chemiluminescent northern blot (right). Analysis is performed in duplicates (R1 and R2). **E)** Utilization of chemiluminescent northern blot for detection of queuosinylated tRNA-His (right) in bacteria using 0.25% APB and TAE based urea-PAA mini gels (left). *Δtgt* serves as non-queuosinylated control of wild-type (wt) strain. Gel and samples were prepared as described in ref. ^6^ **F)** Chemiluminescent northern blot detection of thiolated tRNA-Glu (right) in yeast using 10% urea-PAA mini gels containing 10 μg/mL APM (left). Gel and transfer were performed as described in this protocol.

The method employs a two-column purification strategy using distinct binding conditions. In the first column purification step, the binding conditions (2.3M guanidine thiocyanate (GITC), 35% ethylene glycol diacetate (EGDA)) are optimized for binding RNA molecules longer than 120 nucleotides while shorter RNAs, including tRNAs, remain unbound and are collected in the flowthrough (FT). In the subsequent column purification step, the binding conditions (1.6M GITC, 60% EGDA) are adjusted to capture all RNA species present in the flowthrough from the first column.

***CRITICAL*:** Make sure that all steps are performed at ambient temperature and the samples are mixed thoroughly. Do not place RNA mixed with isolation buffers on ice at any point. Only the initial total RNA dilution and final long and short RNA fractions eluted in RNase-free water should be kept on ice.

***Note:*** tRNA yields depend on the quality and purity of total RNA used for isolation. See Troubleshooting 1 for further guidance.

***Note***: For isolation of long RNAs (>120 nt), perform optional steps a-g.

1. Dissolve and/or dilute 80-100 μg of DNA-free total RNA with RNase-free water to a final volume of 100 μL. **Note:** Total RNA does not need to be DNase treated, but excessive DNA contamination will decrease the tRNA yields. ***Note***: Total RNA should be dissolved in RNase-free water. The ideal concentration is >1 μg/μL. If the concentration of total RNA is <1 μg/μL, concentrate RNA by precipitation or vacuum concentration. Alternatively, perform the first purification step with multiple columns and then pool the FT for the second column step where the short RNAs are bound to the column. For further details, see Troubleshooting 1.
2. Mix the diluted total RNA with 250 μL of Long RNA Binding Buffer (LBB). Vortex thoroughly. ***CRITICAL***: Always use freshly prepared LBB!
3. Add 190 μL of 100% ethylene glycol diacetate (EGDA). Mix thoroughly and incubate for 2 min at ambient temperature. ***Note***: The appearance of a white precipitate is indicative of DNA contamination. This may reduce tRNA yields.
4. To bind long RNAs, load the mixture onto the column (placed into a 2 mL collection tube).
5. Spin column for 1 min at 11 000×*g*.
6. Collect FT. DO NOT discard the FT, as it contains the short RNAs. Proceed to step 7 for isolation of short RNAs. ***Note***: For long RNAs, continue with optional steps a-g at the end of this section.
7. Mix FT from step 6 (∼540 μL) with 675 μL of Short RNA Binding Buffer (SBB). Vortex thoroughly and incubate for 1 min at ambient temperature.
8. Bind short RNAs to silica column:
  i. Load half (∼600 μL) of the mixture to a fresh silica spin column. ***Note***: Maximum loading volume is 700 μL.
  ii. Spin for 1 min at 3 000×*g*.
  iii. Discard FT and reload the column with the remaining solution.
  iv. Spin for 1 min at 3 000×*g*.
  v. Discard FT and continue with step 9.
9. Wash column with 500 μL Chaotropic Wash Buffer (CWB). Spin for 1 min at 13 000×*g*.
10. Wash column with 500 μL Ethanol Wash Buffer (EWB). Spin for 1 min at 13 000×*g*.
11. Repeat step 10.
12. Spin empty column for 2 min 13 000×*g* to dry the column.
13. Transfer column to a new 1.5 mL tube and elute short RNA by adding 50 μL of RNase-free water. Incubate for 1-2 min. Spin column for 1 min at 13 000×*g*.
14. Measure RNA concentration and check purity on a 10% PAA-UREA gel. Store at −80 °C until further analysis.

***Optional***: Isolate long RNA from first column according to the instructions below:

a. Place column to new 2 mL collection tube.
b. Wash column with 500 μL CWB. Spin for 1 min at 13 000×*g*. Discard FT.
c. Wash column with 500 μL EWB. Spin for 1 min at 13 000×*g*. Discard FT.
d. Repeat step c.
e. Spin empty column for 2 min 13 000×*g* to dry the column.
f. Transfer column to new 1.5 mL tube and elute long RNA by adding 50 μL of RNase-free water. Incubate for 1-2 min. Then spin column for 1 min at 13 000×*g*.
g. Store at −80 °C until further analysis.

***Pause point***: The isolated tRNA in RNase-free water can be stored at −80 °C long-term if stored properly and freeze-thaw cycles are avoided.

### Biotinylation of DNA probes

**Timing: 3-4 h**

This section describes a method for enzymatic 3’ end labeling of DNA probes with biotin using Terminal Deoxynucleotidyl Transferase (TdT). The following reaction setup produces 1-3 nt long biotin-16-dCTP tails. Biotinylation efficiency is verified by probe analysis on native PAA gel (Figure 1B).

***CRITICAL***: Before assembling the reaction, allow all components to reach ambient temperature, except for biotin-16-dCTP and TdT enzyme, which should be kept on ice or at −20 °C, respectively.

15. Dilute unlabeled DNA oligo to 1 pmol/μL in RNase-/DNase-free water. Mix well and briefly spin to collect all liquid to the bottom of the tube.
16. Assemble the biotinylation reaction.

**Biotinylation reaction**

| Reagent | Final concentration | Amount per 1 reaction |
| --- | --- | --- |
| RNase/DNase-free water | N/A | 34 μL |
| 10× TdT buffer | 1× | 5 μL |
| 2.5 mM CoCl<sub>2</sub> solution | 0.25 mM | 5 μL |
| 1 pmol/μL DNA probe | 0.1 pmol/μL | 5 μL |
| 1 mM Biotin-16-dCTP | 0.1 mM | 0.5 μL |
| 20 U/μL Terminal Transferase (TdT) | 2 U/μL | 0.5 μL |
| **Total** | **N/A** | **50 μL** |

***CRITICAL***: DO NOT place the individual reaction components or the reaction itself on ice! The use of cold components or cooling down the reaction mixture will result in poor labeling efficiency. For further details see Troubleshooting 2.

***CRITICAL:*** Avoid repeated freeze-thawing Biotin-16-dCTP as this will degrade the nucleotide.

***Note***: When more probe is needed, do not scale up a single reaction; instead, perform multiple reactions.

***CRITICAL:*** TdT may become inactivated if it comes into direct contact with undiluted 10× TdT buffer or CoCl_2_ solution. Hence, when preparing the mastermix for labeling multiple reactions, combine all components except TdT and Biotin-16-dCTP. Mix thoroughly and only then add TdT and Biotin-16-dCTP. Mix again, aliquot 45 µL into tubes and add the DNA probe.

17. Incubate at 37 °C for 60 minutes.
18. Stop the reaction by incubation at 70°C for 10 minutes.
19. Prepare 12% native PAA gel to verify the labeling of probes and let it polymerase for 30-40 min.
20. Prepare gel samples from labeled and unlabeled probe:
  a. 1 μL oligo (1 pmol/μL) + 9 μL MQ + 2 μL 6× DNA LB
  b. 5 μL from finished and deactivated labeling reaction + 1 μL 6× DNA LB
21. Place gel to electrophoresis tank filled with 1×TBE buffer
  a. Remove comb and rinse wells using needle and syringe to remove any polyacrylamide pieces.
22. Load 5 μL from each sample to gel and run at 150 V for 90 min.
23. Disassemble gel electrophoresis and stain gel in 10 000× SYBR Gold dilution in 1×TBE.
  a. Stain the gel in box with gentle shaking for at least 5 min.
24. Visualize gel using UV or blue-light transilluminator.
25. Prepare 10 μL (1 pmol) aliquots of labeled probes.
26. Store aliquoted probes at −80 °C for up to 3 months. ***CRITICAL***: Do not freeze-thaw the probe.

### (t)RNA separation on PAA gels

**Timing: 2-3 h**

In this section, the total RNA or isolated tRNA is separated by denaturing polyacrylamide gel electrophoresis. We routinely use 8–10% gels, but denser (up to 15%) gels can be used for detection of shorter RNAs, specifically tRFs (Figure 1C). The following method can be also used to verify the purity of tRNA fractions (Figure 1A).

27. Thaw tRNA and/or total RNA samples on ice.
28. Prepare a 10% UREA-PAA denaturing gel and allow it to polymerize for 30-40 min.
29. Meanwhile, prepare samples in 2× RNA loading dye. The recommended sample amount is 200 ng/lane for tRNA and 2 μg/lane for total RNA, respectively.
  a. Denature at 80 °C for 5 min.
  b. Place immediately on ice and incubate for 1-2 min. ***Note***: RNA samples mixed with 2× RNA loading dye can be prepared in advance and stored at −80 °C. Before gel loading, denature them as described in step 29.
30. Place gel to electrophoresis tank filled with 0.5× TBE buffer.
  a. Pre-run gel at 200V, 20 min.
  b. Just before loading the gel, rinse wells using syringe and needle to remove urea.
31. Load samples to gel and run at 200 V for 90-120 min. ***Note:*** Electrophoresis time may vary based on the size of the gel (if using a different vertical gel electrophoresis system than described here). For optimal separation, run the gel until the bromophenol blue dye is about to run off the gel.
32. After electrophoresis, disassemble the gel from the apparatus.
  a. Transfer the gel to a RNase-free box containing 10 000× diluted SYBR Gold in 0.5× TBE.
  b. Stain the gel for 3-5 min with gentle shaking.
  c. Visualize the gel by transillumination and save the image (Figure 1C-D, left panel). ***Note***: After imaging, transfer the gel back to the staining solution to prevent the gel from drying out.

### RNA transfer and UV crosslinking

**Timing: 2-3 h**

These steps describe how resolved (t)RNA is transferred from the gel to a positively charged Amersham™ Hybond-N+ nylon membrane. The RNA is then immobilized on the membrane by UV crosslinking. This protocol demonstrates the transfer of ∼14 × 10 cm, 1 mm thick gels, but gels of other sizes can also be used. Where applicable, guidance for handling differently sized gels is provided.

***CRITICAL***: Always use clean tweezers when handling membranes and assembling the blot. When handling the gel, use new and clean nitril gloves.

33. Prepare material for semi-dry transfer
  a. 6 sheets of 3MM filter paper (pre-cut by user to 23 × 14 cm)
  b. Nylon membrane (pre-cut by user to 14 × 10 cm)
  c. PAA gel with our separated (t)RNA in SYBR Gold staining solution or in 0.5× TBE.
34. Soak the filter papers and membrane in 0.5× TBE for at least 20 min.
35. Using tweezers, assemble the semi-dry transfer (from bottom up):
  a. Place the wet 3MM filter paper on the anode (bottom) of the semi-dry transfer apparatus.
  b. Gently roll out the bubbles with a roller.
  c. Repeat steps a-b twice (i.e. 3 sheets of filter paper in total).
  d. Place the wet membrane on top of the filter papers.
  e. Place the gel on the membrane. Align the top left corner of the gel with the membrane. Gently remove bubbles using your fingers. Use clean gloves when handling the gel.
  f. One-by-one, place another set of 3 sheets of wet 3MM filter paper on top of the gel (as described in steps a-b). ***CRITICAL***: Do not use the roller directly on the gel or the membrane. Furthermore, to avoid moving the assembly, only apply gentle pressure when removing bubbles with the roller. Make sure to roll out the air bubbles after each filter paper is added, otherwise it is very easy to trap air bubbles between the layers. The best practice for removing air bubbles is to gently roll from the center of the assembly towards the outer edge in all four directions—center to right, left, up, and down. Rolling from right to left or top to bottom near the edges can reintroduce bubbles, especially in larger gels, where this issue is more pronounced. ***Note***: A short (or shortened) single-use serological pipette can be used instead of a roller.
36. Gently remove excess liquid from the anode using paper towels. ***CRITICAL***: Make sure the stack is sufficiently wetted with 0.5× TBE to ensure uniform transfer.
37. Carefully place the lid on the assembly. Apply even pressure until the lid lock clicks into place.
38. Transfer with 400 mA for 60 min. ***CRITICAL:*** To protect the semi-dry transfer apparatus, set the voltage limit to 24 V when using high current power sources, such as Bio-Rad PowerPac HC. ***Note***: Use 250-300 mA for 60 min when transferring mini gels (8.6 × 6.7 cm).
39. After transfer, gently disassemble the transfer stack with tweezers. ***Note:*** During the disassembly of the transfer stack, the membrane can be marked with a pencil. This is particularly useful when the membrane will be cut into multiple pieces after crosslinking, allowing simultaneous testing of several probes on different sections. ***Optional***: After the transfer, re-stain the gel with 10 000× SYBR Gold solution to ensure that transfer was successful, and no air bubbles were present (Figure 1D, middle panel).
40. Briefly rinse the membrane with RNase-free water to remove any residual gel pieces.
41. Dry the membrane between 2 sheets of 3MM filter paper for few minutes.
42. Before crosslinking, place the dried membrane with the RNA side facing up on a sheet of slightly dampened (with RNase-free water or 2xSSC) 3MM filter paper. Wrap the membrane and filter paper together in single-use plastic wrap or cling film. ***Note:*** The 3 MM filter paper provides support for easier handling and protects the membrane from contamination. When wrapping, avoid trapping any air bubbles between the membrane and the plastic foil. For direct storage of the membrane after crosslinking, use a dry filter paper as support; however, wrapping the membrane in plastic foil may be slightly more difficult in this case.
43. Place the membrane with the RNA side up to the center of the crosslinker. Crosslink twice using UV wavelength 254 nm and 120 mJ/cm^2^ energy as settings. ***Pause point:*** To store the membrane, dry it between two clean filter papers and wrap in plastic foil to protect from scratches and contamination. The UV crosslinked membrane can be stored indefinitely at ambient temperature.
44. Proceed to membrane pre-hybridization.

### Chemiluminescent Northern blot

**Timing: 2 days**

In this step, biotinylated probes specific to tRNAs and tRNA-derived fragments are hybridized to the RNA containing membrane, followed by chemiluminescent probe detection using streptavidin-HRP conjugate and a chemiluminescence substrate. Reagent volumes should be adjusted according to the membrane and container size. Unless otherwise specified, all washes are performed at room temperature.

***CRITICAL:*** Throughout the procedure, always handle the membrane with tweezers to prevent contamination.

45. Warm the pre-/hybridization buffer at 65 °C in a water bath to dissolve any precipitate, mixing occasionally to aid dissolution. ***Note***: Dissolving the precipitate can take up to 60 min. Therefore, start with this step at the beginning of transfer.
46. Once dissolved, transfer 15 mL of hybridization buffer to the hybridization tube and place the tube and the remaining buffer to the hybridization oven (set to 42–45 °C). ***Note:*** The hybridization temperature may need to be individually optimized for each probe. However, 42–45 °C is suitable for most probes designed in accordance with this protocol.
47. Unwrap the crosslinked membrane from the plastic wrap (see step 43) and briefly dip it into a container with membrane wash buffer (2× SSC, 0.1% SDS). Remove excess buffer by lightly tapping one edge of the membrane on a paper towel. ***Note***: Wetting the membrane with membrane wash buffer makes it easier to place the membrane into the hybridization tube.
48. Transfer membrane to pre-warmed hybridization buffer in the tube. ***CRITICAL*:** Ensure that the membrane is correctly positioned, and that the RNA side is facing the inside of the hybridization bottle. To prevent excessive background signal, ensure that the membrane does not dry at any point after adding the hybridization buffer.
49. Incubate the membrane in the hybridization oven for >1 h with slow rotation (10-15 rpm).
50. Next, discard the buffer from the tube and add 15 mL of fresh hybridization buffer.
51. In a 1.5 mL tube, mix 500 μL of hybridization buffer and a 10-40 μL (1-4 pmol) aliquot of biotinylated DNA probe. ***Note***: We successfully used as little as 1 pmol of probe for detection of tRNA isoacceptors and tRNA fragments. However, if the probe is old and/or more samples are loaded onto the gel, a higher probe amount may improve overall signal strength.
52. Denature the probe for 2 min at 95°C, then cool on ice for 2 min. ***CRITICAL***: Do not keep the denatured probe on ice for an extended period, as this can cause the hybridization buffer to precipitate.
53. Add the denatured probe directly to the tube containing the hybridization buffer and membrane. Avoid pipetting the probe directly onto the membrane; instead, dispense it at the bottom of the tube. Mix the tube thoroughly to ensure even distribution.
54. Incubate the membrane in the oven O/N at 42–45 °C with slow rotation (10-15 rpm).
55. On the following day, gently remove the membrane from the hybridization tube and rinse it briefly in membrane wash buffer (2× SSC, 0.1% SDS). ***Note:*** Before removing the membrane from the hybridization tube, make sure that the fresh blocking buffer is ready (see Materials and equipment setup).
56. Wash the membrane in membrane wash buffer (2× SSC, 0.1% SDS) for 5 min.
57. Discard the buffer and repeat the previous step with fresh membrane wash buffer. ***Note***: If the blots have a strong background, more stringent wash buffers can be used (see Troubleshooting 3).
58. Transfer the membrane to a clean container and briefly rinse it with a few mL of PBS wash buffer.
59. Add blocking buffer to completely cover the membrane, then incubate for 60 min with gentle shaking. ***Note:*** Make sure the membrane is completely covered with blocking buffer. 60 mL is usually sufficient for large membranes (14 × 10 cm) while 20–30 mL is adequate for smaller membranes (8.6 × 6.7 cm). However, depending on the size of the container used, additional buffer may be needed. ***Note:*** At this point, take ECL components to warm up at 20–22 °C (if stored at 4 °C).
60. Shortly before finishing the blocking incubation (step 59), prepare a 1:30 000 dilution of streptavidin-HRP conjugate in blocking buffer.
61. Discard blocking buffer from Step 59 and add blocking buffer with streptavidin-HRP (step 60) to the membrane. Incubate 30 min with gentle shaking.
62. Transfer the membrane into a clean container and briefly rinse the membrane with PBS wash buffer.
63. Wash membrane 4 × 5 min with PBS wash buffer with gentle shaking. For efficient washing, the membrane should be freely floating in the buffer.
64. Transfer the membrane into a clean container and briefly rinse the membrane with detection buffer.
65. Add detection buffer to the membrane. Incubate membrane for at least 5 min with gentle shaking. ***Note***: The membrane can be incubated in detection buffer for up to 2-3 hours before chemiluminescent detection. Ensure there is always enough buffer to prevent the membrane from drying.
66. Prepare the chemiluminescent substrate (ECL) by mixing luminol/enhancer solution with stable peroxide solution as indicated by manufacturer. For a 14×10 cm membrane, prepare 5-6 mL of the solution.
67. Remove the membrane from the detection buffer and gently blot the edges on a paper towel to drain any excess liquid.
68. Transfer the membrane to a clean container with the RNA side facing up and add chemiluminescent substrate onto the membrane. Protect from light and incubate for 5 min without shaking. ***CRITICAL***: If the substrate volume is insufficient for the container size, place the membrane RNA side down onto the liquid to prevent it from drying out.
69. Remove the membrane from the substrate and gently blot the edges on a paper towel to drain excess liquid. Do not allow the membrane to dry! Wrap the moist membrane in plastic wrap. Avoid trapping air bubbles between the plastic wrap and the membrane. ***Note:*** *Air bubbles can be gently pressed out with a roller or fingers*.
70. Visualize the signal using a chemiluminescent imaging system (e.g., ChemiDoc MP, Bio-Rad). Optimize the exposure time to ensure efficient signal detection. ***Note:*** For the Chemidoc MP, we recommend using the auto rapid mode with recommended binning size. For weaker signals, manual exposure of 300-1200 s might be necessary. In case no signal is obtained, consult Troubleshooting 4.
71. Take a colorimetric picture of the membrane to see its positioning. Pictures can be overlayed and analyzed in the Image Lab software.

## Results – Expected outcomes

This protocol describes a method for isolating tRNA from yeast and bacteria, but it is readily applicable for other organisms. It effectively enriches for tRNA while longer RNA molecules are efficiently removed. Typical tRNA yields from TRIzol isolated bacterial total RNA^2^ are 9-10 µg, while 4-5 µg can be obtained from yeast total RNA isolated with hot phenol^9^. In contrast, the acidic phenol method^4^ can yield up to 20 µg of yeast tRNA. Isolated tRNA can be utilized for enhanced detection of tRNA-derived fragments (Figure 1A) or further analyzed by sequencing or mass spectrometry.

The Northern blot method described here utilizes enzymatic biotinylation of standard DNA probes (Figure 1B). This allows multiple probe designs to be validated without the need for commercial labeling, which significantly reduces the cost. Furthermore, this increases the sensitivity of the Northern blot, allowing us to distinguish individual tRNA isoacceptors and derived fragments by chemiluminescent detection (Fig. 1C and D). We also demonstrate that this method is compatible with other well-established blotting techniques, such as the detection of queuosinylated or thiolated RNA using 0.25% APB gels (Figure 3E)^10^ or 10 µg/mL APM gels (Fig. 1F)^4^, respectively.

## Limitations

The tRNA isolation method presented here is compatible with virtually any total RNA sample; albeit the main limitation is the minimum amount of total RNA required. For samples with low total RNA input, columns with smaller binding capacities should be used to ensure efficient isolation. Additionally, pre-tRNAs close to or longer than 120 nt will not be detectable in the tRNA fractions.

The success of the Northern blot depends on the design of the DNA probes. Designing probes for tRNA isoacceptors can be challenging due to their high sequence similarity. Furthermore, the obtained signal intensity varies with target tRNA or RNA abundance. Hence, low-abundance molecules may go undetected. It is also important to recognize that Northern blot results are semiquantitative, and complementary methods may be required for a comprehensive quantitative analysis.

## Troubleshooting

### Problem 1

Low yields from tRNA isolation

#### Potential solution

- Too high DNA contamination of total RNA used for isolation. Perform DNase treatment of the total RNA.
- Poor integrity of total RNA. Check RNA integrity by gel electrophoresis or automated electrophoresis system (e.g. Agilent TapeStation, Fragment Analyzer, etc.). If the quality of the total RNA is low, consider using other total RNA isolation methods.
- Increase the amount of total RNA for isolation. We have successfully used up to 250 μg of total RNA with NucleoSpin RNA silica columns.
- Perform the first column purification multiple times (steps 1-6), then pool the tRNA-containing flowthrough and proceed with second column purification (steps 8-14).
- Presence of lipids in the total RNA. Isolate total RNA using protocol for lipid rich samples.
- Check for precipitates in the buffers and prepare fresh buffers, if necessary. Ensure that the EGDA solution did not change color.

### Problem 2

Probe labeling is inefficient or absent

#### Potential solution

- Biotin-16-dCTP was freeze-thawed. Use new aliquot or purchase new compound.
- For some probes, only one biotin-16-dCTP is added. This is normal and does not affect probe functionality.
- TdT enzyme was added to the concentrated reaction buffer during master mix preparation.

### Problem 3

Too high background for Northern blots

#### Potential solution

- Include more stringent washes (e.g. higher buffer temperature or decrease SSC concentration) after step 57.
- Prepare fresh blocking solution.
- Perform longer blocking step with more blocking solution.
- Try new probe design. A low signal intensity will lead to stronger background depending on the exposure time.
- Process the chemiluminescent image using the Image Lab software. Background noise can be reduced by cropping the image to remove excess black space around the membrane, followed by auto contrast adjustment, which enhances the intensity of the signal.

### Problem 4

No chemiluminescent signal

#### Potential solution

- Check biotinylation efficiency of the DNA probe. The probe might have lost the terminal biotin during storage. Analyze the stored probe on a native PAA gel as described in steps 19-24 or perform a dot blot with the probe only. Spot a few probe dilutions onto the membrane and perform detection (steps 55-71).
- Verify the functionality of all buffers by performing a dot blot using a previously validated probe (as described above).
- Verify that transfer was successful by staining the gel post-transfer with SyBr Gold.
- Decrease the hybridization temperature.
- Load more tRNA and/or total RNA onto the gel. This applies for the detection of molecules with low abundance (steps 27-32).
- Try new probe design for same target RNA.
- Make sure the chemiluminescent substrate is not degraded.
- Use a more sensitive chemiluminescent substrate, such as SuperSignal West Pico PLUS, Thermo Scientific, 34580).

## Resource availability

### Lead contact

Further information and requests for resources and reagents should be directed to and will be fulfilled by the lead contact, L. Peter Sarin.

### Technical contact

Technical questions on executing this protocol should be directed to and will be answered by the technical contact, Pavlina Gregorova.

### Materials availability

This study did not generate new unique reagents.

### Data and code availability

This study did not generate new datasets or code.

## Acknowledgments

Authors thank Salla Kalaniemi for technical support and all members of the RNAcious laboratory for further testing of this protocol. This work was supported by the Research Council of Finland (grant no. 354906 to L. P. S.) and the Sigrid Jusélius Foundation (grant no. 230182 to L. P. S.). P. G. is a fellow of the Doctoral Programme in Integrative Life Sciences. The graphical abstract was created with BioRender.com

## Author contributions

Conceptualization, P.G.; experimental methodology, P.G., M-M.H. and M.L; writing, P.G., M-M.H. and L.P.S.; supervision and funding acquisition, L.P.S.

## Declaration of interests

The authors declare no competing interests.

## Notes

### Competing Interest Statement

The authors have declared no competing interest.

